# Water requirement and growth indicators of forest tree species seedlings produced with automated irrigation management

**DOI:** 10.1101/2020.08.24.264879

**Authors:** Mateus Marques Bueno, Paulo Sérgio dos Santos Leles, João Felício Gonçalves Abreu, Jaqueline Jesus Santana dos Santos, Daniel Fonseca de Carvalho

## Abstract

The lack of information regarding the water requirement of tree species promotes water waste in the seedlings production in nurseries. Water requirement, the growth plant factors and water efficiencies for height and diameter were determined for *Schizolobium parahyba* (Vell.) Blake, *Cytharexylum myrianthum* Cham. and *Ceiba speciosa* Ravenna seedlings, under greenhouse conditions and automated irrigation management. We used sewage sludge biosolids as substrate in the seedling phase (280 cm^-3^ tube), and sandy soil material in the initial pot growth phase (18 dm^-3^ pot). In the seedlings phase, four water replacement levels were applied to the substrate, by drip irrigation, meaning average replacement ranging from 40 (V1) to 100% (V4) of species water requirement. Seedlings developed properly and 80 days after emergency, *S. parahyba, C. myrianthum* and *C. speciosa* seedlings received, respectively, 2.40, 1.08 and 0.85 L per plant, for V4. After growth phase (230 DAE), the total water volumes were, respectively, 70.0, 50.3 and 52.7 L per plant. Under adequate water supply, there were rapid recovery and growth of the species, even for the seedlings which showed different height and diameter in the tube phase. The growth plant factors values found were below 0.5 for all species indicating low sensibility to growth, both in height and in diameter, in response to water deficit. Water efficiency indicators point to distinct trends between the two phases, and *C. speciosa* present higher values of water efficiencies for height (80.7 and 17.0 cm L^-1^) and diameter (2.1 and 0.5 mm L^-1^) in both phases.

## INTRODUCTION

Areas requiring some sort of recovery in different degraded ecosystems of Brazil reach about 21 million hectares and are characterized as with legal deficit of native vegetation (Sansevero et al. 2018). Much of this area is located in the Atlantic Forest biome, considered the most altered in Brazil, with an estimate of only 12.5% of its original cover (Santos et al. 2019).

In the formation of stands aimed at forest restoration, it is essential to produce seedlings with high quality and rusticity, which are influenced by the availability of water, both over time and in terms of volume (Keffer et al. 2019), and by the substrate used. Adequate supply of nutrients to seedlings can be performed by using sewage sludge biosolid as substrate (Alonso et al. 2018), as it contains high organic matter content and has appreciable amounts of N and P (Abreu et al. 2017).

Seedling quality is also related to the water supply in the nurseries, which should be carried out in response to the crop water requirement according to the interactions of the environment and water dynamics in the substrate (Keffer et al. 2019). About 49% to 72% of the water applied in reforestation nurseries is lost, varying according to the species (Dumroese et al. 1995). In this context, drip irrigation associated with automated management can promoting adequate water supply for the full development of the crop (Panigrahi et al. 2012), using less water and labor (Dias et al. 2013).

Although there are reports on the growth of native species of the Atlantic Forest tree, most of them do not quantify the water requirement for seedlings, as well as their sensitivity to water deficit, which would contribute to the development of more efficient forest plantations. In order to represent the impact of the use of water resources in the tree seedlings production, we present news growth and water efficiency indicators related to, respectively, water deficit and water volume applied, for the development of seedlings in height and diameter.

The present study aimed to determine the water requirement, the growth plant factors (Gpf) and water efficiencies (WE) of Atlantic Forest tree species, under automated irrigation management in the seedling stage in plastic tube. Also, we evaluated the water requirement and WE during initial growth in pots, in a greenhouse.

## MATERIAL AND METHODS

The experiments were conducted from September 2018 to May 2019 and consisted in the planting of 3 species of native forest tree seedlings, *Schizolobium parahyba* (Vell.) Blake (Guapuruvu), *Cytharexylum myrianthum* Cham. (Pau Viola) and *Ceiba speciosa* Ravenna (Paineira), in 280 cm^3^ plastic tubes (phase 1) and in 18 dm^-3^ pots (phase 2).

The experimental areas are located in the Horticultural Sector of the Agronomy Institute of the Federal Rural University of Rio de Janeiro, municipality of Seropédica – RJ, Brazil (22°45’48” S, 43°41’50” W, altitude of 33 m). The regional climate classified by Köppen’s international system is Aw (Alvares et al. 2013), humid climate and dry winter, with average annual rainfall of 1300 mm and average annual temperature of 22°C.

Seeds from mother trees of the Atlantic Forest of Rio de Janeiro were initially placed in sand boxes. After 20 days of emergency, when the seedlings were about 10 cm tall, their height and collar diameter were measured, and 60 homogeneous seedlings of each species were selected for transplantation to the tubes, which were installed in 4 trays, totaling 15 seedlings per tray. The material was placed on a metal workbench (0.8 × 1.2 × 3.0 m) and one micro-irrigation drip system was installed for each species, with independent flow rate sensors and independent reservoirs, located at 1.70 m above the workbench.

In first phase, from 09/25/1918 to 11/24/2018, the experiments were conducted in a completely randomized design, with 4 treatments (irrigation levels) and 15 replicates (tubes per tray), in climatized environment. The emitters used were spaghetti-type microtubing (Plasnova, mod. PDAEXT001000354), with 0.8 mm diameter and different lengths (0.80, 0.50, 0.35 and 0.20 m), promoting the different irrigation levels. After conducting flow rates tests (Table 1), distribution uniformity coefficients (DUC) greater than 95% were obtained.

**Table 1.** Mean flow rate of the emitters (L h^-1^) in the respective treatments, for the tree forest species

| Treatments | Experiments/Species |  |  |
| --- | --- | --- | --- |
|  | *S. parahyba* | *C. myrianthum* | *C. speciosa* |
|  | Mean flow rate of the emitters (L h<sup>-1</sup>) |  |  |
| V1 | 1.2 | 1.1 | 0.7 |
| V2 | 1.8 | 1.5 | 1.1 |
| V3 | 2.1 | 2.3 | 1.4 |
| V4 | 2.9 | 2.6 | 1.8 |

The substrate used in the tubes was pure biosolid, obtained from a sewage treatment plant managed by the Rio de Janeiro State Water and Sewage Company (*Companhia Estadual de Águas e Esgotos* – CEDAE). Physical characteristics indicate the existence of large porous space (0.70 cm^3^ cm^-3^), low bulk density (0.74 g cm^-1^) and approximately 70% of the average particle diameter between 1.0 and 0.5 mm. The physical-hydraulic parameters of the substrate were obtained using the simplified evaporation method (Schindler 1980), through the commercial device Hyprop® (Pertassek 2015), and indicated low water holding capacity. This substrate has values of N (1.61%), P (0.68%), K (0.27%) and organic carbon (9.66%), which provide the mineral supply without the need for fertilization, especially in the initial growth stage of the seedlings.

Water management was realized by simplified irrigation controller (SIC) (Medici et al. 2010), which operates in response to soil water tension and is regulated by the level difference between a porous capsule (sensor) and a pressure switch. This device has been used in several field (Mello et al. 2018) and greenhouse studies (Gomes et al. 2017), using different soil or substrates for plants. It is presented as an alternative in irrigation management (Bezerra et al. 2019) and has the advantage of being built with low cost materials (Valença et al. 2018). Two actuators were used for each plant species studied, independently installed in two tubes with treatment corresponding to the highest flow rate (V4), totaling 6 actuators for the 3 experiments.

The volume of water applied per treatment over time was measured by water flow sensors (mod. YF-S201b), connected to an Arduino MEGA board (mod. 2560), responsible for data storage. In order to avoid any percolation losses, a system composed of a rain sensor (mod) with direct actuation by relay was installed. Two sensors were installed in each experiment, just below the tube that received the largest volume of water.

At 60 days after start of irrigation monitoring, when the seedlings reached average heights of 36.4 cm (*S. parahyba*), 25.9 cm (*C. myrianthum*) and 32.4 cm (*C. speciosa*), 16 seedlings of each species (4 of each treatment) were transferred to 18 dm^-3^ pots filled with soil material collected from the 0-40 cm layer of a *Quartzpsamments*, located in the municipality of Seropédica, RJ, Brazil. This soil had A horizon about 20 cm deep, followed by C horizon up to 120 cm of drilling. The dominant particle-size fraction was sand and loamy sand, the mean pH was 5.2 and base saturation (V) in the A horizon was equal to 36%, indicating low fertility.

Using the same experimental design, this second phase, totaling 48 seedlings, was conducted in a plastic greenhouse, simulating the field conditions, except for the water supply, performed by irrigation. In a period of 150 days, from 11/24/2018 to 04/24/2019, a uniform water depth was applied for all plants of each species by the independent irrigation system whose management was also carried out by the AAI, with sensors installed in the seedlings of V4 treatment. The irrigation system consisted of one dripper per pot (mod. PCJ - Netafim), with nominal flow rate of 2.0 L h^-1^, and the water volume applied was measured by previously calibrated hidrometer (Alpha mnf/FAE) installed on the main line. Uniformity tests indicated DUC above 97.0%. At 150 days after emergency, *C. myrianthum* plants required fertilization, because they had visual symptoms of nutritional deficiency.

The meteorological monitoring of the experiments was carried out by a WatchDog 2000 Series weather station (Spectrum Technologies, Illinois/USA), containing sensors of temperature, relative humidity, solar radiation and wind speed, and with data storage every 30 minutes. The average daily temperature during the first phase was 26.6 °C, with little variation due to the climatized environment. The average relative humidity air was around 80% and the average solar radiation was 8.1 MJ m^-2^ dia^-1^. In second phase, the average daily temperature was 27.4 °C, with average amplitude of 10.8 °C. The average relative humidity air was around 44.0% and the average total radiation was 21.0 MJ m^-2^ d^-1^.

In both phases, nondestructive growth analyses for plant height (cm, distance from plant collar to apical bud, with a graduated ruler) and collar diameter (mm, with a digital caliper) were performed every 15 days, in first phase, and every 30 days, in second phase.

To verify whether the premises of the analysis of variance were met, the normality and homogeneity of the residuals were assessed by the Bartlett *test*, at the level of 5% probability. Meeting these criteria, the analysis of variance was performed using the t test, with a significance level of 5%. After rejecting the null hypothesis, polynomial regression analyzes were performed for the accumulated volume applied and irrigation regime factors. All statistical analyzes in this study were performed using software package R, version 3.6.0.

Using the same concepts of the yield response factor (Ky) (Garg and Dadhich 2014), the growth plant factor related to water deficit for the development of seedlings in height and diameter (Gpf) was calculated by Eq. 1.

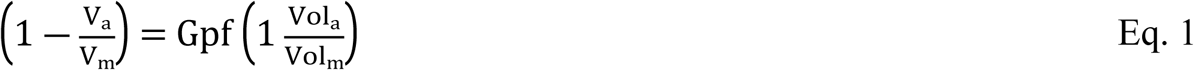

where V_a_ is actual variable value (height - cm, or diameter - mm), V_m_ is maximum variable value (height - cm, or diameter - mm), Vol_a_ is actual volume applied (L) and Vol_m_ is maximum volume applied (L).

Since this relationship above is linear, Gpf correspond to the slope of the regression line, which was obtained interactively by maximizing the V_m_ value (height or diameter) so that the equation intercept with the ordinate axis became equal to zero. The procedure was performed in an electronic datasheet (MS Excel™), using the Solver module (Carvalho et al. 2016). The interpretation of Gpf was done according to Doorenbos and Kassam (1979), for Ky.

Water efficiency (WE) based on height (HWE) and diameter (DWE) was obtained according to Eq. 2.

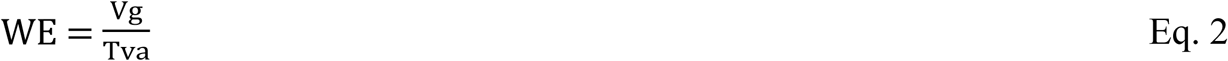

where Vg is the variable growth (height - cm, and diameter - mm) and Tva is the total volume applied, in L per plant.

## RESULTS AND DISCUSSION

At 60 days after start of irrigation monitoring, from 20 to 80 days after emergency (first phase), the species reached on average the minimum standard for field planting in all treatments, which corresponds to the range from 20 to 40 cm for height and diameter greater than 3 mm (Souza Junior and Brancalion 2016), for quality seedlings produced in 280 cm^3^ tubes. During this period, the irrigation systems were actuated 73, 43 and 52 times for the *S. parahyba, C. myrianthum* and *C. speciosa* species, respectively, applying total volumes of 0.91, 1.49, 1.74 and 2.40 L per plant, 0.46, 0.62, 0.95 and 1.08 L per plant, and 0.33, 0.52, 0.66 and 0.85 L per plant, for V1, V2, V3 and V4, respectively. The irrigation system was turned on more than once a day for the three species in 14, 5 and 10 days (2 activations), and in 6, 3 and 4 days (3 activations), respectively, for the *S. parahyba, C. myrianthum* and *C. speciosa* species. In addition to meeting the seedlings water requirement, the large number of drives of the SIC is associated with the low water holding capacity in the substrate.

There were positive responses of height (H) and collar diameter (D) as a function of the increase on water volumes applied to all species seedlings (Fig1).

**Fig 1.**
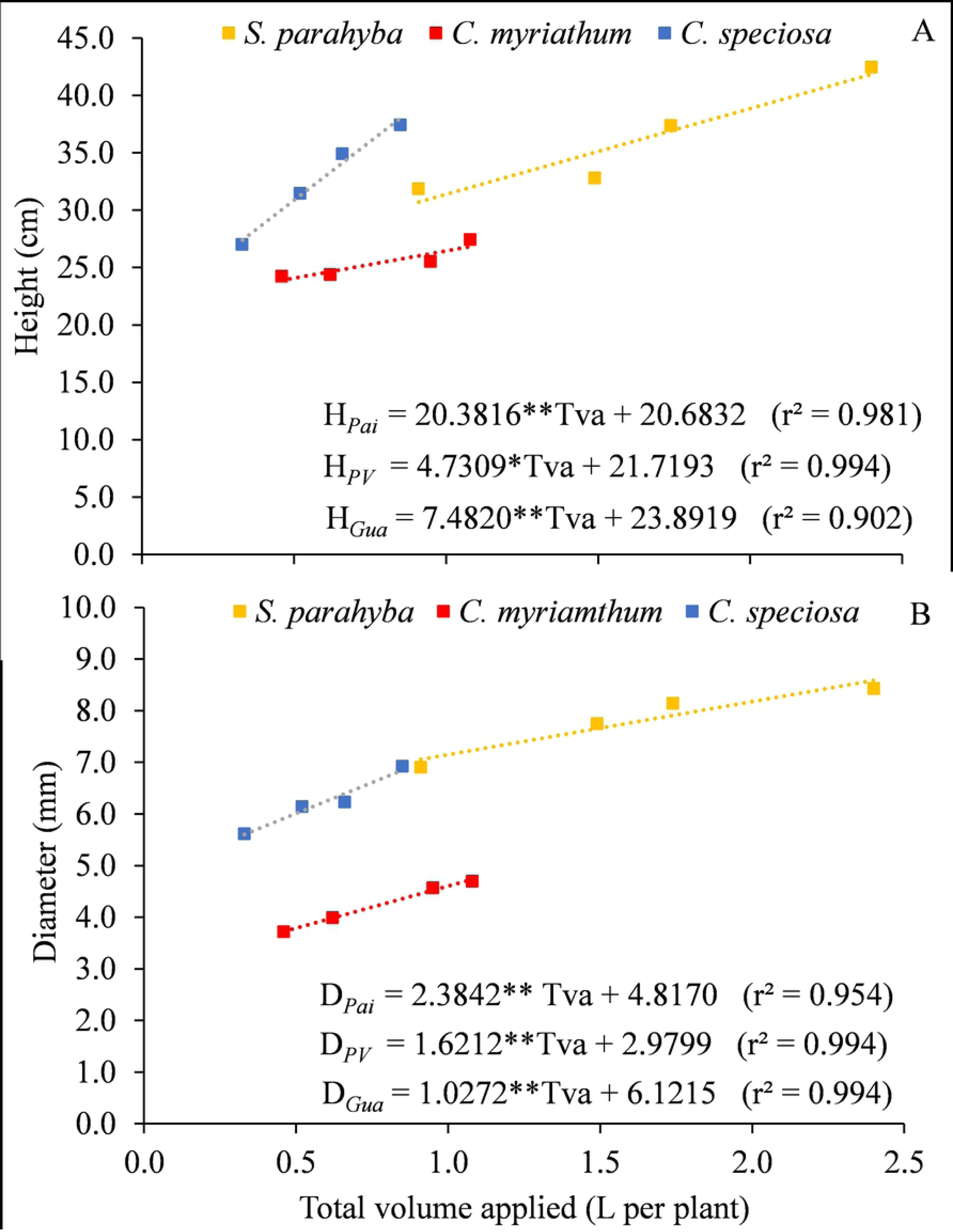
Variations of height (A) and diameter (B) as a function of the total volume applied (Tva) on treatments for three forest species, in the tube phase. *p<0.05, **p<0.01

It is worth pointing out that the SIC used in this study provides water for plants in response to their development (Medici et al. 2010), maintaining the moisture corresponding to the water capacity of the substrate. Thus, the volumes of 2.40, 1.08 and 0.85 L per plant, for the V4 treatment, corresponding to the water requirement for the *S. parahyba, C. myrianthum* and *C. speciosa* species, respectively.

The variations in height (Fig1A) and diameter (Fig1B) of the species were proportional to the different volumes applied, justifying the linear adjustment of the regression models. The lowest angular coefficients indicate lower responses to seedling growth and were obtained for *C. myrianthum* (4.7309 cm L^-1^ – height) and *S. parahyba* (1.6212 mm L^-1^ – diameter). *C. speciosa* species had the highest growth rates in response to the amount of water applied, with values of 20.3816 cm L^-1^ in height (Fig1A) and 2.3842 mm L^-1^ in diameter (Fig1B). The amount of water demanded by *S. parahyba* seedlings was, on average, 171.1% higher than the volume applied to the *C. speciosa* seedlings, although the differences in height (10.7%) and diameter (25.4%) were proportionally smaller. The difference in the relative water demand for these two species probably occurred due to their phytoecological regions of occurrence. According to Carvalho (2003), *S. parahyba* occurs in the alluvial plain and the beginning of the slopes, while *C. speciosa* frequently occurs in different environments, such as in some areas of the caatinga domain. *C. myrianthum* had the lowest growth in height and diameter, reaching just 27.43 cm and 4.69 mm, respectively. As it is a non-pioneer species, its lower growth is explained by the lack of shading precisely in the initial stages of development (Morais Júnior et al. 2019).

Calculated from Fig1, the maximum height/diameter ratio (H/D) values for *S. parahyba* (5.0), *C. speciosa* (6.5) and *C. myrianthum*. (5.6) are within the range below 8.0 indicated by Souza Junior and Brancalion (2016). This relation is used to assess the quality of forest seedlings, because, in addition to reflecting the accumulation of reserves, it ensures greater resistance and better fixation in the soil (Artur et al. 2007). The occurrence of water deficit in the substrates tends to limit the growth of tree seedlings in terms of diameter and height, as well as modify the ideal H/D ratio for a given species (Duarte et al. 2016). However, the variations in the H/D ratio observed in this study were not enough to provide signs of stiolation in the seedlings. According to Monteiro et al. (2016), the effects of irrigation on the height and diameter of tree or fruit species seedlings vary depending on the plant, size, growth habit, leaf type and size, substrate characteristics and cultivation container, besides the microclimatic conditions of the growing environment.

The growth of the seedlings in relation to the water volumes applied per plant throughout the first phase in the V4 treatment is presented in terms of height (Fig2A) and diameter (Fig2B) for three species. *S. parahyba* species reached greater height (42.5 cm) and diameter (8.4 mm), followed by *C. speciosa* (37.4 cm, 6.9 mm) and *C. myrianthum* (27.4 cm, 4.7 mm). Using a substrate composed of organic matter, peat and vermiculite, in field conditions, *S. parahyba* seedlings reached a maximum height of 36.8 cm, with daily wetting carried out manually (Caron et al. 2010).

**Fig 2.**
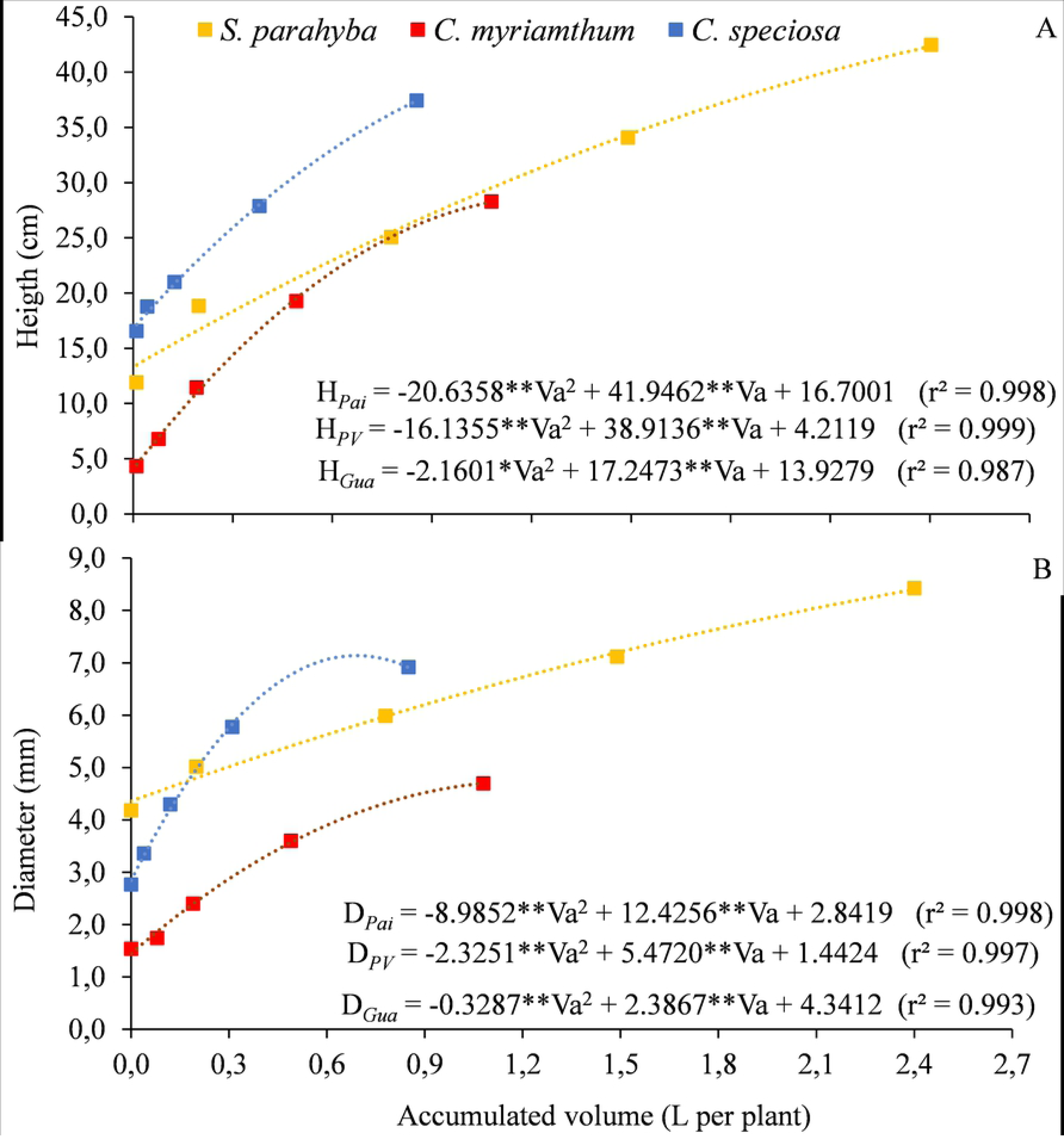
Variation of height (A) and diameter (B) as a function of the water volume applied throughout the experiment (Va) for three forest species, in the treatment with 100% water replacement (V4) for the tube phase. *p<0.05, **p<0.01

In the regression models, the low coefficients of the quadratic terms for *S. parahyba* species mean a trend of linearity in the growth of seedlings in height (Fig2A) and diameter (Fig2B), especially after the second analysis, held on 10/10/2018, indicating a more uniform development throughout the evaluation period. *C. myrianthum* species started the development phase in tubes with the smallest height (4.3 cm) and diameter (1.5 mm) and showed an intermediate growth rate. The highest growth rate in height (Fig2A) and diameter (Fig2B) was found for *C. speciosa* species, despite the lower total water volume applied (0.85 L per plant).

The growth of the species in terms of height and diameter increased proportionally with the dates of evaluation, consequently demanding greater volumes of water. From the first to the second (09/25 to 10/10) and from the fourth to the fifth growth analysis (11/09 to 11/24), the seedlings of *S. parahyba, C. myrianthum* and *C. speciosa* grew, respectively, 6.9 and 8.4 cm, 2.5 and 8.1 cm, and 2.2 and 9.6 cm in height (Fig2A). In terms of diameter, the variations were 0.8 and 1.3 mm, 0.2 and 1.1 mm, and 0.6 and 1.1 mm, respectively (Fig2B). In these periods, the applied volumes were 0.02 and 0.91 L per plant, 0.08 and 0.59 L per plant, and 0.04 and 0.54 L per plant, respectively. In addition, from 09/11 to 11/24, the SIC was turned on 31 (42.5%), 24 (55.8%) and 29 (55.8%) times for the *S. parahyba, C. myrianthum* and *C. speciosa* species, respectively, applying 0.910 (37.9%), 0.5907 (54.6%) and 0.540 (63.5%) L per plant. These values indicate the importance of an adequate irrigation management in order to meet the seedlings water requirements, which varies over time.

Under pot conditions, simulating cultivation in the field, the water volumes applied were 70.0, 50.3 and 52.7 L per plant, respectively, for the *S. parahyba, C. myrianthum* and *C. speciosa* species, with 182, 150 and 171 actuations of the irrigation systems. The irrigation systems were turned on more than once a day in 29, 18 and 29 (2 activations), and in 28, 22 and 24 days (3 activations), respectively, for the *S. parahyba, C. myrianthum* and *C. speciosa* species.

The heights and diameters of the species did not differ statistically at 5% significance level, by Tukey test, over the 5 months of evaluation, indicating that under the condition of adequate water supply there is rapid recovery and growth of the species, even for those which showed different diameter and height in the tube phase (Fig1). Thus, through the automatic management of the irrigation system by SIC, the supply of water to plants was adequate and guaranteed full development of the seedlings. However, in field conditions, this situation may not occur due to increases in temperature and the intensity and frequency of drought periods caused by climate change (Contin et al. 2014). The growth of the seedlings in terms of height (Fig3A) and diameter (Fig3B) indicate a greater water use for *S. parahyba* species, however *C. speciosa* reached greater height (123.5 cm) and diameter (27.3 mm). The amount of water applied to *C. myrianthum* species was similar to *C. speciosa*, although the seedlings reached, on average, 94.5 cm in height and 14.2 mm in diameter.

**Fig 3.**
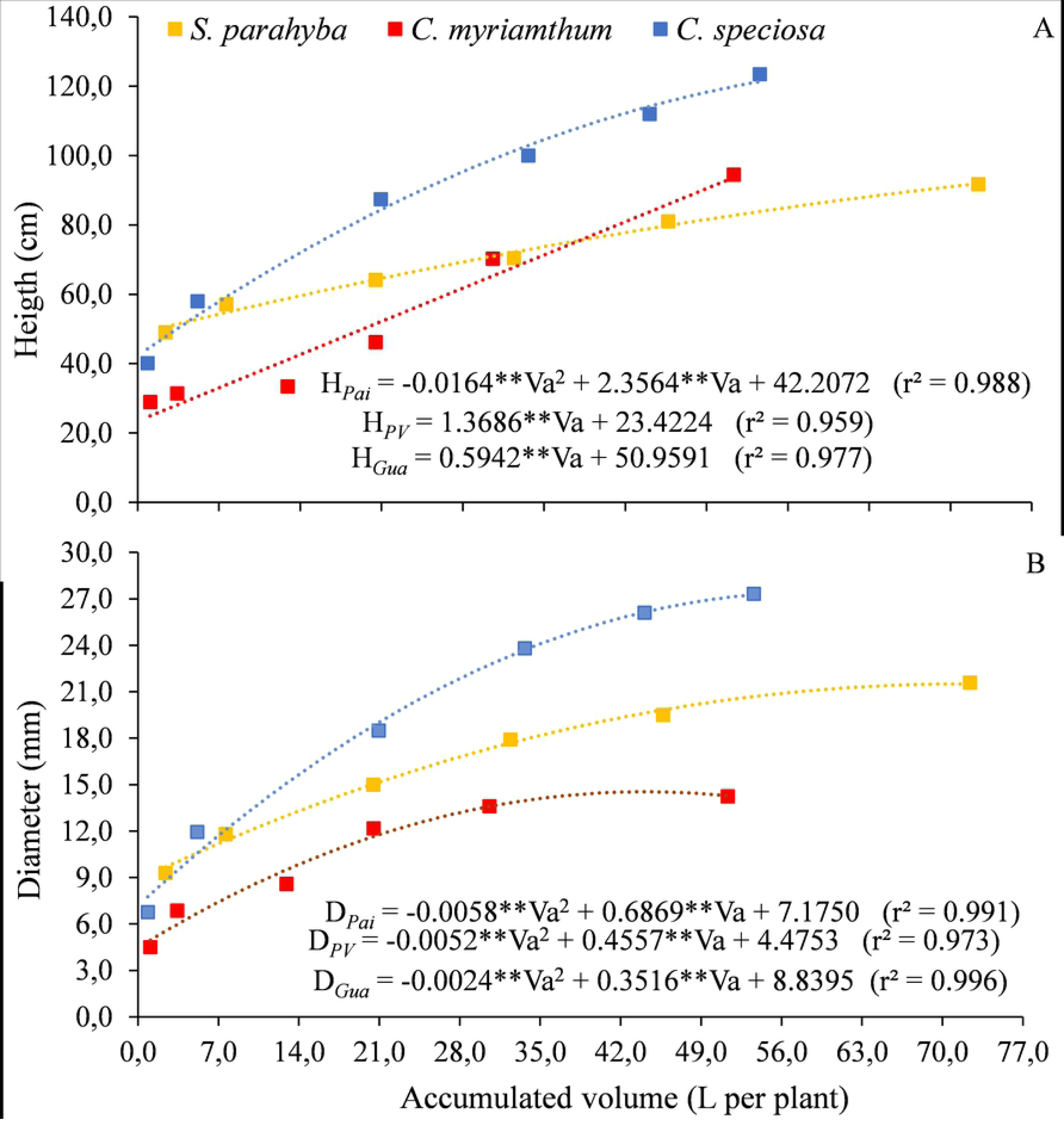
Variation of height (A) and diameter (B) as a function of the water volume applied throughout the experiment (Va) for three forest species, in the pots phase. **p<0.01

The S. parahyba species presented the same trend observed in the tube growth phase, with a lower rate compared to the other species and a uniform growth between the evaluation dates (Fig3A). From the first to the second (11/24/2018 to 12/24/2018) and from the fifth to sixth growth analysis (03/23/2019 to 04/24/2019), the seedlings grew 8.1 and 10.8 cm, respectively, with applied volumes of 5.24 and 26.7 L per plant. In these periods, the average height and volume applied to C. myrianthum seedlings varied, respectively, 2.5 and 24.3 cm, and 2.3 and 20.8 L per plant.

These species presented a linear growth trend in height, while the *C. speciosa* species presented a second order polynomial tendency, despite the low coefficient of the quadratic term (−0.0164) (Fig3A). Unlike the previous ones, *C. speciosa* seedlings had higher growth in height (29.38 cm) and applied volume (15.78 L per plant) between the second (12/24/2018) and third (01/23/2019) measurements, when the irrigation system was turned on 51 times. In this period, the highest incidence of radiation (24.7 MJ m^-2^ d^-1^) and the maximum temperature (40.4 °C) of the experiment were recorded.

The greater volume of water applied to *S. parahyba* species among the fifth to sixth growth analysis (26.7 L per plant) was sufficient to promote plant growth in 2.10 mm in diameter, while for *C. myrianthum*, the volume of 20.75 L per plant provided variation in diameter of just 0.6 mm, in the same period (Fig3B). For these species, the irrigation systems turned on 54 and 43 times, respectively. The volume of 15.78 L per seedling of *C. speciosa*, applied between the second and third measurements provided a variation in the diameter of 6.6 mm. There was greater proportional growth in diameter than in height for the species *S. parahyba* and *C. speciosa*, while for *C. myrianthum* this growth was similar.

Morais Júnior et al. (2019) conducted a comparative study between pioneer and non-pioneer species of the Atlantic Forest and concluded that *C. speciosa* and *S. parahyba* are among those with the highest growth rates, receiving a score 8 and 9, respectively, in a classification from 0 to 10, to compose projects of recovery of degraded areas. Souza et al. (2020) evaluated native forest species in mixed plantations in Brazilian Savanna at 6.4 years old and the *C. speciosa* was among the highest growth, presenting average stocks of volume and biomass of 114.03 m^3^ ha^-1^ and 52.99 Mg ha^-1^, respectively.

*S. parahyba* is a fast-growing tree (up to 45 m^3^ ha^-1^ yr^-1^) and recommended and widely cultivated as ornamental or for restoration purposes (Abreu et al. 2014), and not being very demanding on soil fertility, which facilitates its adaptation in different locations site (Morais Junior et al. 2019). For 13 years, Schwartz et al. (2017) evaluated the development of an area of regeneration with *S. parahyba* in the Amazon region. The species showed a volume increase of 3.1 m^3^ ha^-1^ year^-1^ for individuals with diameter greater than 25 cm and more than 30% of the planted seeds were able to germinate, establish and grow until reaching a diameter greater than 25 cm. This field result combined with the present study indicates that *S. parahyba* has high capacity for water use and, consequently, for forming stands.

The Gpf values found were below 0.5 for all species, as a function of both height and diameter (Fig4), indicating low sensitivity to water deficit, despite showing rapid growth.

**Fig 4.**
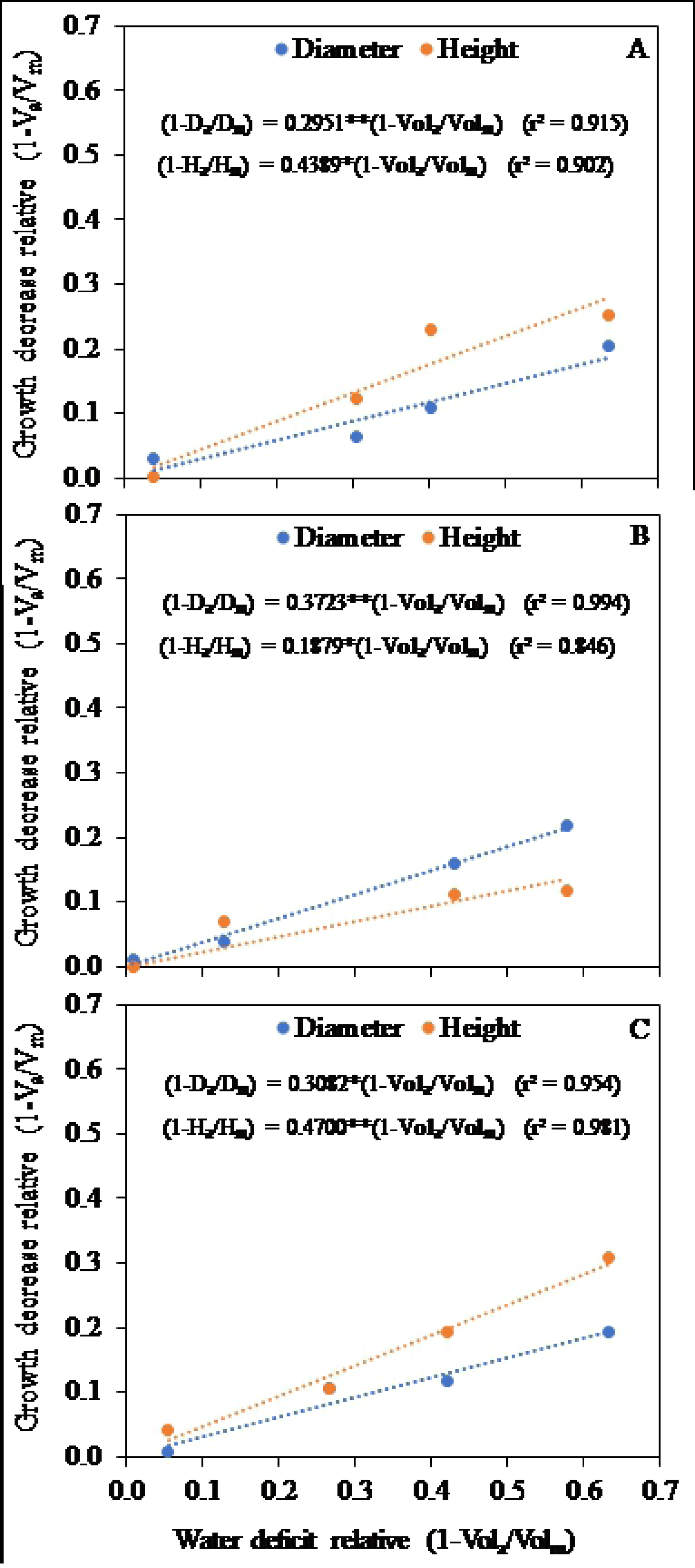
Relationship between growth decrease relative, in height and diameter, and water deficit for *S. parahyba* (A), *C. myrianthum* (B) and *C. speciosa* (C) seedlings. *p<0.05, **p<0.01.

This trend is confirmed by the results obtained in phase 2 (pots) when seedlings of all species from the different treatments in phase 1 (tubes) had satisfactory development, showing rapid recovery and growth under adequate water supply. The lowest and highest Gpf values (Fig4) obtained refer to the height variable for the *C. myrianthum* (0.1879) and *C. speciosa* (0.47) species, and they are related to the respective slopes observed in the response functions of Fig1A.

We understand that the interpretation of the proposed index (Gpf) helps in the management of seedling production systems, however it cannot be compared numerically with other results in the literature as it is not normally used to evaluate the sensitivity of forest seedlings to water deficit. In any case, Keffer et al. (2019) state that the low sensitivity of forest tree species to water deficit in the seedling phase occurs due to having only one vegetative stage. It can be affirmed that the reductions in height and diameter are relatively small compared to the evaluated levels of water deficit, indicating that the species have high capacity for forming stands in initial processes of regeneration, even under conditions of low rainfall, provided that there is minimal water supply in the soil.

Water efficiency (WE) indicators point to distinct trends between the two phases (tube and pot), when the values are compared as a function of height (HWE) and diameter (DWE) (Table 2). In general, WE increase with the reduction in the volume applied for tube phase, and higher values are obtained for *C. speciosa*, followed by *C. myrianthum* and *S. parahyba*. In the pot phase, the differences between the values are reduced due to the recovery observed in seedlings from the treatments with deficits applied in phase 1. Except for *S. parahyba* in treatment V4, all HWE values obtained are statistically equal for the respective species.

**Table 2.** Water efficiency indicators related to total volume applied for the development of seedlings in height (HWE) and diameter (DWE), for *S. parahyba, C. myrianthum* and *C. speciosa* species

|  |  | Water efficiency indicators |  |  |  |
| --- | --- | --- | --- | --- | --- |
| Species | Treatment | HWE (cm L<sup>-1</sup>) |  | DWE (mm L<sup>-1</sup>) |  |
|  |  | tube | pot | tube | pot |
| *S. parahyba* | V1 | 35.018 a* | 1.072 ab | 7.582 a | 0.296 a |
|  | V2 | 22.013 b | 0.986 b | 5.196 b | 0.297 a |
|  | V3 | 21.475 cd | 0.993 b | 4.676 bc | 0.283 a |
|  | V4 | 17.694 d | 1.267 a | 3.510 c | 0.298 a |
| *C. myrianthum* | V1 | 52.676 a | 1.760 a | 8.074 a | 0.257 a |
|  | V2 | 39.343 b | 2.010 a | 6.431 b | 0.300 a |
|  | V3 | 26.877 c | 1.909 a | 4.808 c | 0.318 a |
|  | V4 | 25.401 c | 1.841 a | 4.345 c | 0.277 a |
| *C. speciosa* | V1 | 80.653 a | 2.055 a | 17,002 a | 0.469 a |
|  | V2 | 60.513 b | 2.052 a | 11.809 b | 0.477 a |
|  | V3 | 52.930 bc | 2.098 a | 9.433 c | 0.482 a |
|  | V4 | 44.039 c | 2.305 a | 8.138 d | 0.510 a |

\*Means followed by the same letter in the column do not differ by Tukey test at 5% probability level, for the same species.

Although the best seedlings development has been observed under 100% water suppression, the field monitoring and water efficiency values showed that there may be a good development even in seedlings that received less water. In absolute terms, *C. speciosa* was the species that had the highest values of water efficiency, indicating greater capacity to use this resource. It must be pointed out, however, that all species showed satisfactory growth and were efficient in the use of water, thus being recommended for processes of regeneration of native forests. Tree planting contributes to a rapid recovery of the forest structure, thus providing an adequate habitat for the restoration of the ecological succession (Holl and Aide 2011).

## CONCLUSIONS

*S. parahyba, C. myrianthum* and *C. speciosa* species exhibit greater growth when subjected to automated irrigation management, at a level of 100% water supply, when planted in substrates composed of pure biosolid.

Growth plant factors (Gpf) for the seedling stage are lower than 0.5, indicating low sensibility to growth in the face of water deficit.

The best level of water efficiency was found for the *C. speciosa*, indicating that this specie has greater capacity to use this resource.

The studied species present rapid growth and are efficient in the water use, which implies a good development in conditions of hydric disponibility variable, common in the reforestation processes.

## ACKNOWLEDGMENTS

This research was funded by Fundação de Amparo à Pesquisa do Estado do Rio de Janeiro (FAPERJ) grant number E-26/202.909/2018) and Coordenação de Aperfeiçoamento de Pessoal de Nível Superior - Brasil (CAPES) - grant number 001.

We thank the Federal Rural University of Rio de Janeiro, specifically the PPGA-CS, GPASSA and LAPER.

